# Common marmosets (*Callithrix jacchus*) excel in a one-trial spatial memory test, yet perform poorly in a classical memory task

**DOI:** 10.1101/2025.02.05.636629

**Authors:** Sandro Sehner, Flávia Mobili, Erik P. Willems, Judith Burkart

## Abstract

When quantifying animal cognition, memory represents one of the most tested domains and is key to understanding cognitive evolution. Memory tests thus play an important role in comparative cognitive research, yet slight variations in the experimental settings can substantially change the outcome, questioning whether different memory tests tap into different memory systems or whether they test memory at all. Here, we first assessed memory performance of 16 common marmosets (*Callithrix jacchus*) in two distinct paradigms varying in their format and delay. First, we examined marmoset memory in a 24-hour delay memory test (24h-DMT) in which they could freely explore an environment with three novel objects of which one contained food. We examined their retention the day after, and the procedure was iterated cumulatively with previous objects remaining in the enclosure until the marmosets had to choose the correct out of 30 objects. Second, we administered a classical delayed response test (DRT) in the same animals with three objects and a maximum delay of 30 seconds. In the DRT, marmoset performance was poor and not better than chance after 15 seconds already. However, individuals excelled in the 24h-DMT, performing above chance level after 24h even with tenfold the number of objects to choose from compared to the DRT. Moreover, individual performances in the two tests were not correlated, and typical age effects on memory could not be detected in both experiments. Together, these results suggest that the two tests explore different domains, and that the 24h-DMT examines long-term memory. The outcome of the DRT is more difficult to assign to memory since individuals performed only moderately even in the 0-second delay condition. This puts into question whether this task design indeed tests memory or other cognitive processes.

## INTRODUCTION

Animal cognition is often tested using standardized cognitive tests. For instance, the primate test battery (Banerjee et al., 2009; Herrmann et al., 2007) uses a set of various standardized tests to examine cognitive abilities across domains and species. One of these abilities is memory, which has been assessed in a variety of species. Memory tasks can vary in the way that they are performed, e.g. with regard to the number of choices, or the delays (a few seconds to a couple of minutes) imposed before an animal is allowed to make a decision. Overall, individuals perform worse with an increasing delay in contexts where the maximum delay is only a couple of minutes (Amici et al., 2010; Darusman et al., 2014; Flaim & Blaisdell, 2023; ManyPrimates et al., 2022; Reynoso-Cruz et al., 2021; Schubiger et al., 2016). Larger-brained primates such as apes usually outperform the smaller-brained ones and in particular do so in longer delay conditions (Amici et al., 2010; van Schaik et al., 2024). However, even if an animal does not make the correct choice, the information about a location of a hidden food reward is not necessarily always lost but merely may be temporarily inaccessible (Martin-Ordas & Call, 2011). For instance, chimpanzees (*Pan troglodytes*), bonobos (*Pan paniscus*) and orangutans (*Pongo pygmaeus*) showed a performance decline from a delay of two minutes up to four hours. However, afterward, performance increased again, and with a delay of 24 hours, it was on par again with the delay of two minutes. Overall, performances in memory tests can be influenced by a variety of intrinsic and extrinsic factors. In the following, we will explore three factors that may influence performance in memory tasks: test format, cognitive decline associated with natural aging, and attention allocated to the cognitive test.

First, the format of the test can affect an individual’s performance. The oldest and most common way to test memory in animals is through the classical DRT (Hunter, 1913). The experimenter hides food in one of several cups in a way that the animal can see the hiding process and the individual has to choose one of the cups after a predefined delay. Only when the individual chooses the cup where the food is hidden will it receive the reward. This test has been widely applied across primates (e.g., Call 2001; Barth and Call 2006; Amici et al. 2010; Martin-Ordas and Call 2011; Schubiger et al. 2016). Although this paradigm minimizes extraneous variables, thus fostering comparability between study sites (ManyPrimates et al., 2022), the classical DRT presents an unrealistic scenario in which monkeys generally perform poorly compared to their great ape relatives and commonly appear to forget hiding locations within less than a minute (Amici et al., 2010; ManyPrimates et al., 2022; Schubiger et al., 2016). Furthermore, recently it has been suggested that the classical DRT does not measure memory but rather taps into the domain of attention (van Schaik et al., 2024). However, slight adjustments to the test format can improve performance. For instance, common marmosets (*Callithrix jacchus*) and squirrel monkeys (*Saimiri sciureus*) performed poorly in a two-choice DRT without a clear trajectory of performing worse in conditions with a longer delay (Schubiger et al., 2016). Contrary to what one would expect, increasing task difficulty by decreasing the chance level of making a correct choice (nine cups instead of two) increased overall performance and revealed a typical forgetting curve with decreasing performance with increasing delays.

Another way to assess memory is in open foraging paradigms, simulating a more natural setting for the animals. These, “naturalistic” test scenarios provide evidence for impressive abilities to memorize food locations, even in small-brained species that usually perform poorly in the DRT (ManyPrimates et al., 2022; Menzel & Juno, 1985). In a series of experiments, saddle-back tamarins (*Saguinus fuscicollis*) could freely explore a testing enclosure that contained a set of baited and unbaited objects. The individuals could learn within one trial about the location and properties of an object that contained food and would revisit these objects after a delay of 24 hours significantly earlier and faster than objects that previously did not contain food. In contrast to the classical DRT, this delayed memory task (24h-DMT) showed that these small-brained tamarins could memorize food locations for an extended period after a single learning trial. Similar effects have been found in marmoset monkeys when assessing spatial memory in open-foraging tasks (Abreu et al., 2020; Easton et al., 2003; MacDonald et al., 1994; Platt et al., 1996; Vannuchi et al., 2020). In contrast to the DRT, such open foraging paradigms provide animals with both visual and spatial clues, which could improve performance. In fact, studies have shown that performances in tasks requiring spatial memory decline faster with age than performances in tasks relying on non-spatial memory (Barnes et al., 1987), which may be due to aging effects on the hippocampus (reviewed in Rosenzweig & Barnes, 2003). Therefore, when relying solely on spatial memory, performance is expected to be lower compared to scenarios that utilize both spatial and visual memory Furthermore, animals can explore cues for a longer time period than in the DRT, which may further consolidate memory. Hence, it is expected that animals perform better in open-foraging paradigms than in the classical DRT. However, it remains unknown whether individual performance in the 24h-DMT correlates with that in the classical DRT, which would be expected if both tasks indeed measure memory.

Second, it is widely accepted that many cognitive abilities decline with aging (Salthouse, 2019), and memory is no exception. In humans, memory performance constantly increases until it reaches its peak at around 20 years of age after which it declines linearly (Brockmole & Logie, 2013). Similar age effects on memory have also been investigated quite extensively in animals (reviewed in Berchtold & Cotman, 2009; Gallagher & Rapp, 1997). One explanation for the decrease in cognitive performance is the so-called *neural dedifferentiation* in which neural processes become less selective and cognitive processes like memorizing information are impaired (Alexander et al., 2012; Berchtold & Cotman, 2009; Koen et al., 2020). However, although *neural dedifferentiation* predicts this cognitive decline, findings from animal studies have yielded inconsistent results in this regard. For instance, whereas performance in pigeons in an associative learning task declined with age, memory was unaffected (Flaim & Blaisdell, 2023). In a study with long-tailed macaques (*Macaca fascicularis*), age did predict memory performance in the delayed response task (Darusman et al., 2014). In contrast, in a recent comparison across 41 primate species, age improved memory performance overall (ManyPrimates et al., 2022), which may be due to a lack of older individuals and a bias toward younger individuals with still increasing cognitive performance in this study. Alternatively, it may be that the conducted tests effectively did not examine memory, but instead measured a different cognitive component altogether that is not affected by increasing age.

Third, a component that will influence the result of memory tasks, or in fact any cognitive test, is attention toward the task at hand. Attention has been closely linked with cognitive performance in animals and humans (Decker & Duncan, 2020; Schubiger et al., 2020). For instance, in great apes, it has been shown that establishing signals and securing the attention of the tested animals can increase cognitive performances (Mulcahy & Call, 2009; Mulcahy & Suddendorf, 2011). In another example, marmosets paid little attention when the costs of being unattentive were rather low, but increased their attentiveness when chances of being correct decreased (Schubiger et al., 2016). Especially in these smaller species where individuals inherently face a higher predation risk, individuals might pay more attention to their surroundings than to the actual test, which if not controlled for, can impair their cognitive performance (Schubiger et al., 2015).

In this study, we tested common marmosets in two different tasks examining memory performance. First, we tested individuals in an experimental design that resembles the 24-hour DMT (Menzel & Juno, 1985), followed by examining memory using the classical DRT. Marmosets are group-living, arboreal, and relatively small-brained platyrrhines (Solomon & Rosa, 2014; Yamamoto et al., 2010) that show high levels of vigilance also in captivity (Brügger et al., 2023; Phaniraj et al., 2024). Their natural distribution ranges from the dry caatinga to the humid Atlantic forest (Garber et al., 2019). Marmosets are generalists feeding on a wide range of food items (Souto et al., 2007) for which they use a variety of foraging techniques. Generalists are thought to exhibit a superior spatial memory compared to specialists (Henke-von der Malsburg et al., 2020), and indeed wild common marmosets optimized their foraging route in a spatial memory task based on single-trial learning (Abreu et al., 2020; Xavier et al., 2024). Performance in spatial memory was observed over short-term (4.5 hours) and long-term (11 days) in which marmosets approached previous food locations more often compared to locations that did not contain food. Hence, like the tamarins in the 24-hour DMT (Menzel & Juno, 1985), memory performance in a more natural setup surpassed the usual outcome of the classical DRT. However, a second crucial difference to the classical DRT is that in both studies (Abreu et al., 2020; Menzel & Juno, 1985) individuals were tested within their social unit rather than alone. Not only can isolation manifest in increased stress (Ward & Webster, 2016) impairing cognitive performance, but being in a group can have positive effects on cognitive performances of problem-solving (Sehner et al., 2022), which are not restricted to, but may well include, finding food locations. Hence, the question remains whether the testing performance was better than expected since animals were not tested individually or because the experiment itself resembled a more natural setup.

This study aimed at examining the ecological and content validity, as well as the commensurability, of two commonly used tests. Therefore, we employed the 24h-DMT and the classical DRT in common marmosets. We predicted that animals would excel in the more realistic setup of the 24h-DMT, whereas performance in the classical DRT would be moderate at best and rapidly decrease with increasing latency. We furthermore explored which factors contributed to individual performance in both tasks. Following the general assumption that memory performance decreases with age, we expected that older individuals would do worse in both experiments if they indeed measured memory (Darusman et al., 2014). In addition, we explored whether sex could influence performance. Since previous studies have pointed out that males react stronger towards experimenters than females and are less attentive (Schubiger et al 2015), we expected a better performance of females in the DRT but not in the 24h-DMT, given that the latter was conducted without an experimenter present. Lastly, assuming that both tests examine memory we expected an intraindividual correlation between test performance scores.

## Methods

### Subjects & housing

In this study, we tested 16 (9 females) adult common marmosets (*Callithrix jacchus*) from five captive family groups. The age range of the individuals was from 1.5 to 18 years (for details see Table S1 in Online Resource 1). Older animals have experiences with cognitive tasks, whereas it was the first cognitive experiment for the three individuals under three years of age. However, the only individuals who have experience with memory tasks (the DRT) are Kapi and Vito. All animals were born in captivity and housed in heated indoor enclosures with *ad libitum* access to outdoor enclosures (if weather conditions allowed it). Both, indoor and outdoor enclosures were enriched with natural groundcovers and natural climbing structures. In addition, all enclosures included additional hiding opportunities as well as access to heating lamps. Animals were fed several times a day with a vitamin-enriched porridge in the morning, a mixture of fruits and boiled vegetables over midday, and a snack composed of either animal protein (e.g. fish or boiled egg) or gum arabic. Animals had *ad libitum* access to water and were never food or water-deprived.

### Experiment 1: Twenty-four hours delay memory task (24h-DMT)

#### Procedure

To measure memory over a 24-hour time delay we adapted Menzel and Juno’s (1985) experimental approach and introduced the animals to a novel environment (an indoor enclosure 2.5m height x 1.8m width x 3.5m depth) similar to their home enclosure, henceforth called “testing enclosure”. To habituate the animals to the testing enclosure, we allowed the whole family group to explore the novel environment for one hour on the two days prior to the experiment. The testing enclosure was equipped with several metallic and plastic climbing structures that allowed the animals to reach virtually any point in the enclosure. We did not use any wood structures as these would be difficult to clean between sessions (which is vital, see below). During the experiment and habituation, water was available *ad libitum* from water dispensers. After the habituation phase, individuals had access to this area only during test sessions. During both the habituation and the experiment, we attached two GoPro Hero 7 White at the front-and back of the enclosure. The whole testing enclosure was cleaned with warm water after each family group was tested; to prevent distinct groups’ scent-marking interference (Lazaro-Perea et al., 1999). Due to the cleaning process, only two families could be tested per day. One family was tested in the morning after the porridge feeding but before the fruit/vegetable feeding, whereas the second family was tested in the afternoon after the fruit/vegetable feeding but before the afternoon snack.

The experiment was composed of 10 sessions, with each session comprising two trials that were conducted on two consecutive days. Whether animals were tested in the morning or afternoon was randomized between sessions, but remained consistent within a session to ensure a 24-hour gap. On the first trial of each session, three new test objects were added to the enclosure, accumulating to 30 objects in total in the final session (see Fig. S1 in Online Resource 1). The objects were composed of 10 small plastic bottles adorned in different styles and thus differed in colors, patterns, and surface materials, 10 Lego^TM^ containers differing in size, shape, and colors, and 10 different household objects. Each set of three novel objects for a new session contained one of the customized containers, Lego^TM^ pieces, and household objects. One of the newly added objects was randomly selected to be baited, whereas the other two new objects were never baited throughout the entire experiment. Note, that we took care to balance the number of baited objects from each type over the 10 sessions. The bait was a mixture of one-fourth of nut cookie, two raisins, two pine nuts, and a pinch of gum arabic for each animal. The three novel objects used on each trial were left in a glass container closed with the bait used that day for at least 15 minutes before the experiment started. Hence we controlled for the possibility that animals would approach the baited novel object because it would simply smell the novel food, since all novel objects should at least have some food odor on them. The baited object contained food on both trials of each session. When moving on to the next session, all objects from previous sessions remained in the testing enclosure but none of them was baited again. Hence, objects could have three “baiting states”: baited, previously baited in an earlier session, and never baited. For instance, during the first session, both at the first and second trials, there were three objects only, with only one containing food. In the second session, both on the first and second trials, there were six objects in total, three old ones and three new ones. One of the new ones was baited, and the other five were empty, including the previously baited object from session 1 (Fig. 1). The number of objects in the testing enclosure increased cumulatively as the experiment progressed, reaching 30 objects in the final session (1 object that was baited, nine that were previously baited, and 20 never baited). We randomly assigned the locations of all objects prior to the experiment and used identical locations for all families. Once a location was assigned to an object this location remained the same across subsequent trials and sessions. All individuals were tested separately for 15 minutes and after each trial, the experimenter checked whether the animal had found the food. The order in which the animals within a family were tested was random and depended on the motivation of the individuals. Animals could enter a system of tubes attached to their home enclosure and walk to the test enclosure. The tube system ended in the small cage inside the test enclosure. After an animal entered the small cage the tube system was blocked for the duration of the experiment. To enter the test enclosure, the experimenter lifted a small guillotine door, which was lowered once the animal entered the test enclosure. After the session, the experimenter lifted the guillotine door again and the animal could return to its respective home enclosure using the tube system. Before the next animal was tested, we cleaned food leftovers and rebaited the object if necessary.

**Fig. 1.**
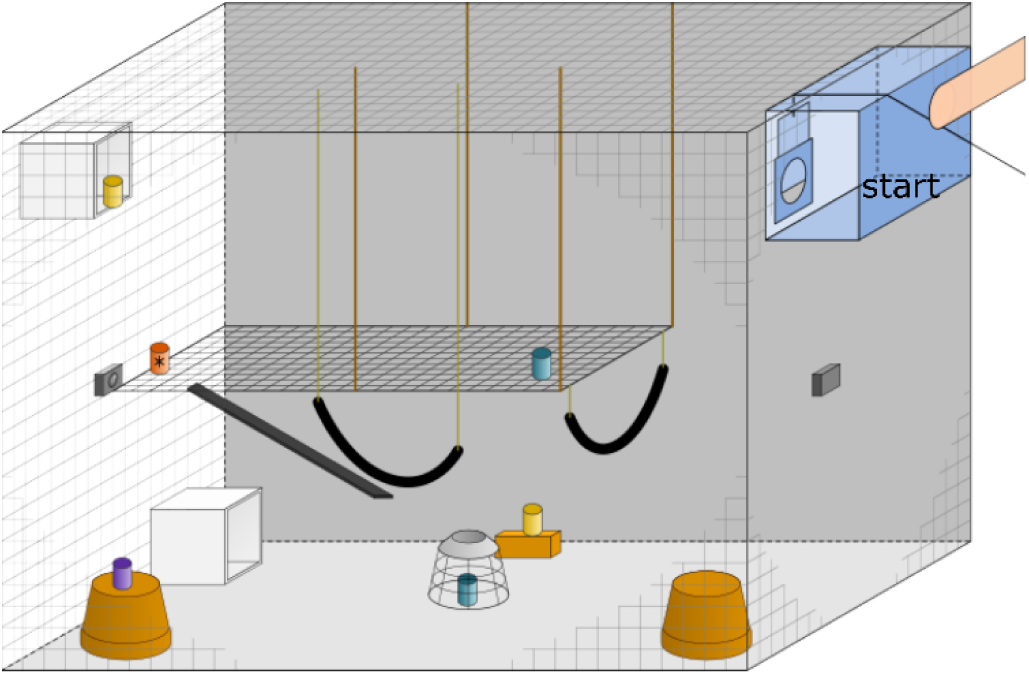
Schematic drawing of the testing enclosure. The small cage inside the enclosure was made of a grid, and the blue pattern differentiates it from the enclosure’s grid. Note, that for visual clarity not all of the 30 used objects are depicted. The small cups represent the objects during the second session. The blue objects are never baited objects from previous sessions; the purple object represents a previously baited object from an earlier session, the yellow objects are the novel never baited objects; the red container represents the baited object of this session. Cameras are indicated with the small black boxes at the front and the back of the enclosure. See the online article for the color version of this figure.

#### Data scoring

All trials were video recorded with two GoPro Hero 7 White and analyzed with INTERACT software V18.1.4.4 from MANGOLD GmbH. The two cameras were positioned on opposite sides of the enclosure (Fig. 1), to monitor whole test enclosure. Each trial started on the preceding frame when the animal started to move away from the guillotine door and entered the test enclosure, and lasted for 15 minutes. We coded the order in which the animal interacted with each object. We also noted the latency for the first interaction with a currently-baited object. An interaction was defined as touching a test object or sniffing/watching the object from a distance up to one arm-length. The coding system consisted of two levels. First, we coded which object was interacted with and then we coded the baiting state of that object of that particular session. A choice was counted as correct choice if an animal interacted first with the baited object before it approached any other object. A second observer coded 20% of all second trials determining when and with which object an individual interacted first. Inter-observer reliability was assessed using Cohen’s Kappa, which yielded a value of 0.83, indicating almost perfect agreement between the two observers.

### Experiment 2: Classical delayed response task (DRT)

After the 24h-DMT, we tested the same 16 individuals using the classical DRT (see also ManyPrimates et al., 2022). Animals were tested individually in a compound (80 cm height x 40 cm width x 40 cm depth), inside their home enclosure. The compound (henceforth “test compound”) was attached to the inside of the home enclosure front grid. During periods between experimental sessions, animals had *ad libitum* access to the test compound. Thus, animals could habituate to the test compound during the entire time unless another individual was tested. To avoid interference, the experimenter moved the group members who were not tested through a tube system to an adjacent room where they had food, water, and enrichment items.

The general setup included a white table suspended by two metal hooks in front of the test compound and outside the home enclosure (Fig. 2). The white table was affixed to the enclosure only during test trials and was removed once a session was completed. Attached to the top of the table was a wooden board (40 cm width x 40 cm depth), and three white opaque cylinder-shaped cups (5 cm high x 3 cm in diameter) were placed on top of the board. The cups were fixed with the opening downwards to maintain the same even distance throughout the test. The board was attached to a pair of drawer tracks and could be moved out of the subject’s reach. The experimenter stood behind the board and turned the three cups with the opening facing the subject. Next, the experimenter showed the bait to the subject and placed it in front of one of the cups. The position could be (1) left, (2) middle, or (3) right – from the experimenter’s perspective. After that, the experimenter erects the cups from left to right, thereby hiding the food item. The delay conditions were short (0-second delay), medium (15-second delay), or long (30-second delay). After the respective delay, the experimenter slid the board towards the subjects while avoiding giving any clues about the hiding position. The marmoset could then choose by pointing toward or touching one of the cups through the grid. If the individual made an ambiguous choice, the trial was not counted and repeated. An ambiguous choice was considered when an animal reached in between two cups and/or did not touch any cup. The trial was also repeated if the subject did not choose within one minute. After the marmoset chose a cup, the board was pulled back, and the chosen cup was lifted and laid down. If the cup revealed the bait, the subject received it as a reward. If the cup did not contain the bait, the subject did not receive a reward, and the same food item was used in the subsequent trial. After revealing the chosen cup, the experimenter lifted the other cups, and a new trial was started.

**Fig. 2.**
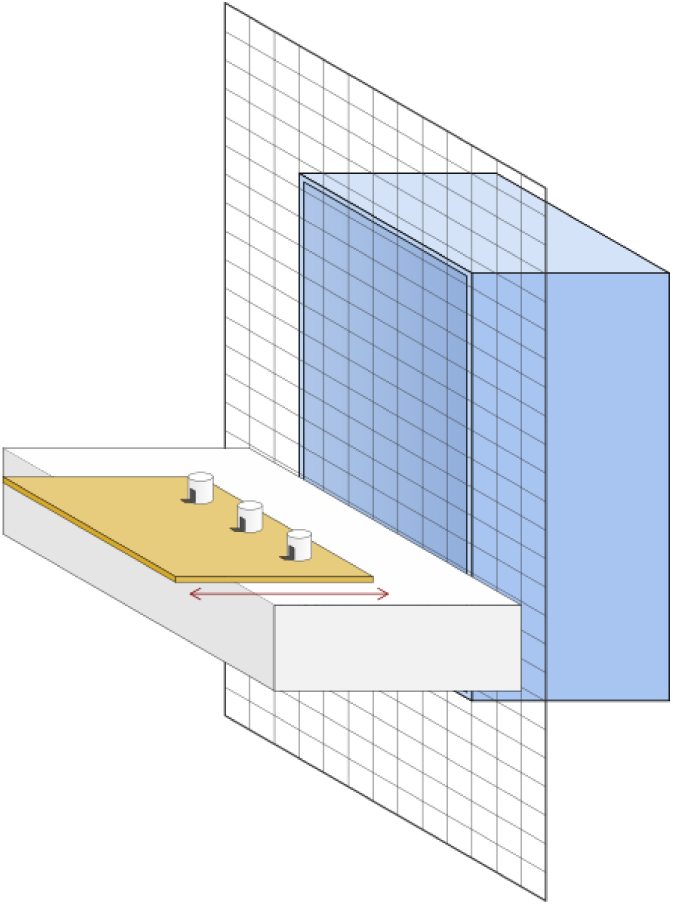
Schematic drawing of the test compound. The front grid represents the enclosure grid, and the test compound inside the enclosure was made of a grid, and the blue pattern differentiates it from the enclosure’s grid. The wooden table can be moved back and forth. The cups were held with a small stripe of tape so as not to fall and interfere with the experiment. See the online article for the color version of this figure.

#### Training

Before starting the test sessions, each subject was trained to ensure reliable choice behaviors. This procedure was not required by the ManyPrimates et al. (2022) project but advised, as the animals had to consistently point to, or reach for, visible food items placed on the table board before the start of the experiments. Accordingly, subjects went through a pre-test phase in which the experimenter followed the same procedure but did not impose any time delay. After the subject reached the criterion (≥ 70% correct choices within 12 consecutive trials) it entered the test phase on the next day. The duration of training varied between subjects, but all 16 individuals passed the criterion of making the correct choice in 70% or more in 12 consecutive trials without delay. Correct choices were those were the animal choose the baited container. In contrast to the ManyPrimates et al. (2022) project, the subject received one warm-up trial without delay at the beginning of each test day. This trial was not considered and served only to attract the subject’s attention as well as to probe its motivation to participate on a given day. Two predefined criteria for stopping a session were used: (1) the subject did not make a choice in one minute in three consecutive trials, or (2) the subject was no longer attentive to the table and/or emotionally aroused and indicated it wanted to leave the test compartment. If the subject showed at least one of these criteria, we stopped the session immediately and restarted the trial the following day.

The ManyPrimates et al. (2022) project provided the hiding location and delay conditions (see Table S2 in Online Resource 1). The hiding position was randomized across trials with the following constrictions: the exact location could not occur more than two consecutive times, and each hiding location occurred an equal number of times per delay. Trials were grouped in blocks, with each block comprising three trials of the same delay (short, medium, and long). The number of trials per session varied among individuals, with the constraint that there were at least three trials per test day. If less than three trials were finished, the session was terminated, and trials were discarded and repeated the following day.

#### Data scoring

All trials were video recorded with a GoPro Hero 9 Black, scored live using ManyPrimates et al. (2022) check sheets, and confirmed later with the video. We coded with which cup a subject interacted first after the board slid forward. We also coded the position of the indicated cup (left, middle, or right). If the indicated cup hid the food, the response was coded as correct. A second person scored 20% of all trials scoring the choice of an animal. The inter-observer reliability was assessed using Cohen’s Kappa, which yielded a Kappa value of 0.93, indicating almost perfect agreement between the two observers. This result was statistically significant (z = 13.7, p < 0.001).

#### Statistical analysis

All statistical analyses were performed in R-studio (R Core Team, 2020). To test whether animals remember the location of the last baited object in the 24h-DMT, we ran a binomial Generalized Linear Mixed-effects Model (GLMM) with a log link-function. The dependent variable scored whether the first interaction was with the currently baited object or not (yes/no). The model did not include fixed effects, however, *individual identity* nested in *family identity*, and *session number* as random effects. We also included the probability of making a correct choice by chance as an offset term. The offset was defined as the log of 1 over n, where n was the total number of objects in each session, and thus varied from 1/3 (first session) to 1/30 (last session). In addition, we conducted a Monte Carlo simulation generating 10.000 random choices for each session amounting to 100.000 random choices for all sessions. Again, we adjusted the probability of making a correct choice by chance level from 1/3 in the first session, to 1/30 in the last session. We furthermore matched the number of animals in the simulation to the empirically observed number of animals that found the baited object in our experiment in the first session. For instance, whereas for the first session, we simulated 10.000 random choices by 16 animals, the last session was based on 10.000 random choices by 11 animals. Thus, the subsequent comparison of the simulated data to the sum of observed first interactions with baited objects on second trials is much more objective.

We formulated a Cox proportional hazards mixed-effects model for the *latency of the first interaction* with the baited object to examine whether animals would find the baited object faster in their respective second trials. The model included the *individual identity* nested *in family identity* and *session number* as random factors, and *trial* as a predictor variable.

To examine whether individual characteristics affected performance, we calculated a binomial GLMM including *age*, *sex,* and *status* as predictor variables, and *session number* and *individual identity nested in family identity* as random effects. Our response variable was structured as a two-column matrix: one column indicated the *number of correct choices* and the other indicated the *number of failures*.

To test whether animals would remember the location of the hidden food item in the classical DRT, we ran a binomial GLMM with a log-link function. First, we examined how individuals performed over all trials, regardless of the delay. The model did not include fixed effects but the family identity as a random effect. We also included the probability of making a correct choice by chance (1/3) as an offset term. In a subsequent analysis, we tested whether the *delay condition* affected the performance. For this, we set the *delay* as a contrast and included it as a predictor in the model. Lastly, we tested whether animals were better than expected by chance for each *delay condition*, calculating a binomial GLMM with the same structure as used for the overall performance.

Similar to the first experiment, we examined whether individual characteristics affected performance, we calculated a binomial GLMM including *age*, *sex,* and *status* as predictor variables and *family identity* as random effect. Our response variable was structured as a two-column matrix: one column indicated the *number of correct choices* and the other indicated the *number of failures*.

## Results

Overall, we recorded 320 trials, equally divided into 160 trials of first and second trials over all sessions. In 80% of all the trials the individual found the baited object, resulting in 256 successful trials (see Table S1 in Online resource 1).

We examined whether animals would remember the location after one trial learning and switch from previously baited objects in previous sessions to novel objects that are baited (Fig. 3). We performed an analysis restricting correct choices to currently baited objects only. The intercept was significantly positive (GLMM: estimate = 2.55; standard error = 0.47, z-value = 5.46: *p* < 0.001), indicating that individuals interacted more frequently first with the currently baited object (73 out of 129 trials) than would be expected by chance alone. We also calculated the number of correct choices in each session to control for difficulty, since the increasing number of objects may have increased the difficulty level in each following session. Our results demonstrated no increase or decrease in the number of correct choices throughout the experiment (see Table S3 in the Online Resource 1 for the table of polynomial contrasts).

**Fig. 3.**
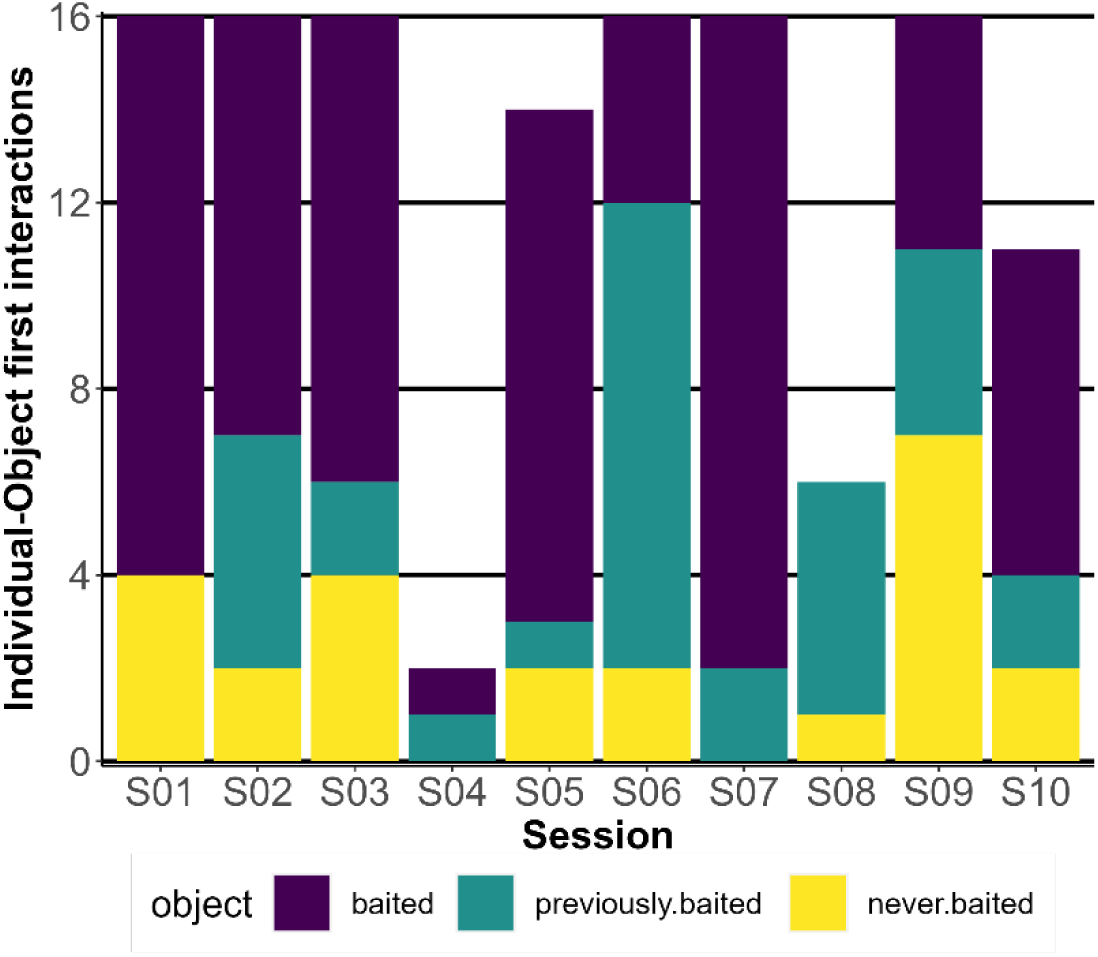
The number of individuals who found the food in the second trial of each session, separated by their first interaction with a baited object (purple), a previously baited object (turqoise), or a never baited object (yellow) in their respective second trial. See the online article for the color version of this figure.

We further examined whether the observed behavioral pattern was likely the result of individual memory and one-trial learning or random choice. We compared the random choices extracted from the Monte-Carlo simulation with the observed first choices of a baited object (Fig. 4). This analysis thus did not consider previously baited objects. In every session, except for sessions four and eight, the observed correct choices were significantly higher than the simulated ones (Table 1). On average, the observed value is 2.15 ± 1.42 times bigger than the 97.5% theoretical quantile.

**Fig. 4.**
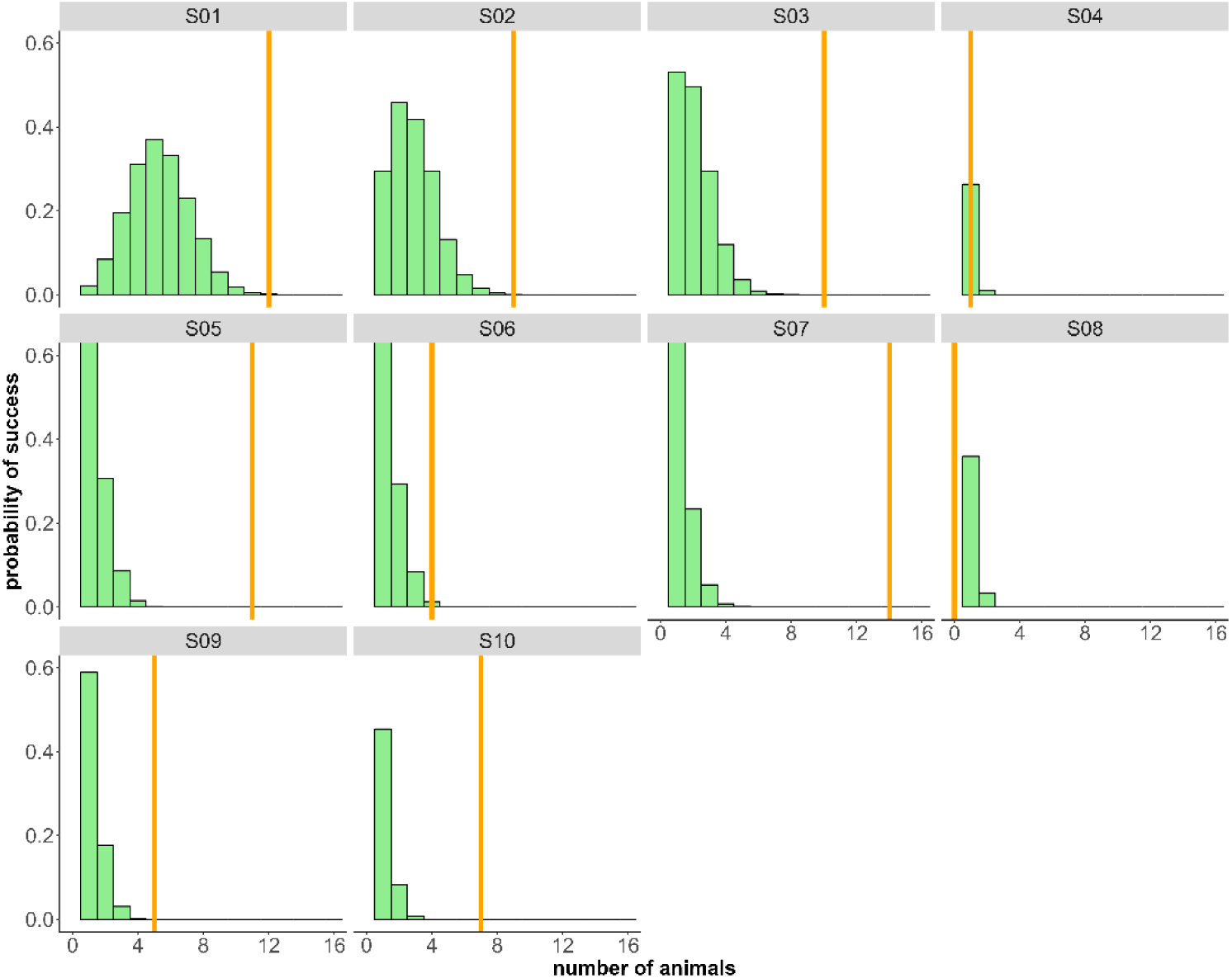
Simulated probability of the number of animals interacting with the correct object (currently baited object) based on random interactions (green histogram). The simulated data for each session is based on the number of animals that found the correct object in the first trial. The orange bar represents the observed number of animals interacting first with the correct object in the second trial. In eight out of ten sessions, a significantly higher number of animals than expected by chance interacted first with the currently baited object (see also Table 1). See the online article for the color version of this figure.

**Table 1.** 95% quantiles and the median of the simulated distribution of the number of animals that would interact correctly by chance per session. Observed number of animals that first interacted with the baited object (N) and the p-value compared to the simulated distribution. Bold values represent < 0.05. Considered are those trials 2 for each session where the individual found the bait during trial 1.

| Session | Number of considered trials | 2.50% | 50% | 97.25% | N | p-value |
| --- | --- | --- | --- | --- | --- | --- |
| S01 | 16 | 2 | 5 | 9 | 12 | **< 0.001** |
| S02 | 16 | 0 | 3 | 6 | 9 | **< 0.001** |
| S03 | 16 | 0 | 2 | 4 | 10 | **< 0.001** |
| S04 | 2 | 0 | 0 | 1 | 1 | 0.15 |
| S05 | 14 | 0 | 1 | 3 | 11 | **< 0.001** |
| S06 | 16 | 0 | 1 | 3 | 4 | **< 0.01** |
| S07 | 16 | 0 | 1 | 3 | 14 | < 0.001 |
| S08 | 6 | 0 | 0 | 1 | 0 | 1 |
| S09 | 16 | 0 | 0 | 2 | 5 | < 0.001 |
| S10 | 11 | 0 | 0 | 2 | 7 | < 0.001 |

In the next step, we examined whether animals would find the correct object faster in the second trial than in the first trial of a session. Hence, we compared the latencies of finding the correct object in the first trials with the latencies of finding the correct object in the respective second trials. A Cox mixed effect model revealed that an animal found the baited object earlier during its second trial than on its first trial (proportional hazard ratio = 1.91, CI_95_% = 1.46 to 2.49, p < 0.001). In other words, at any given instant in time, an individual was 1.91 times more likely to already have found the correct object in the second trial than in the first trial (Fig. 5).

**Fig. 5.**
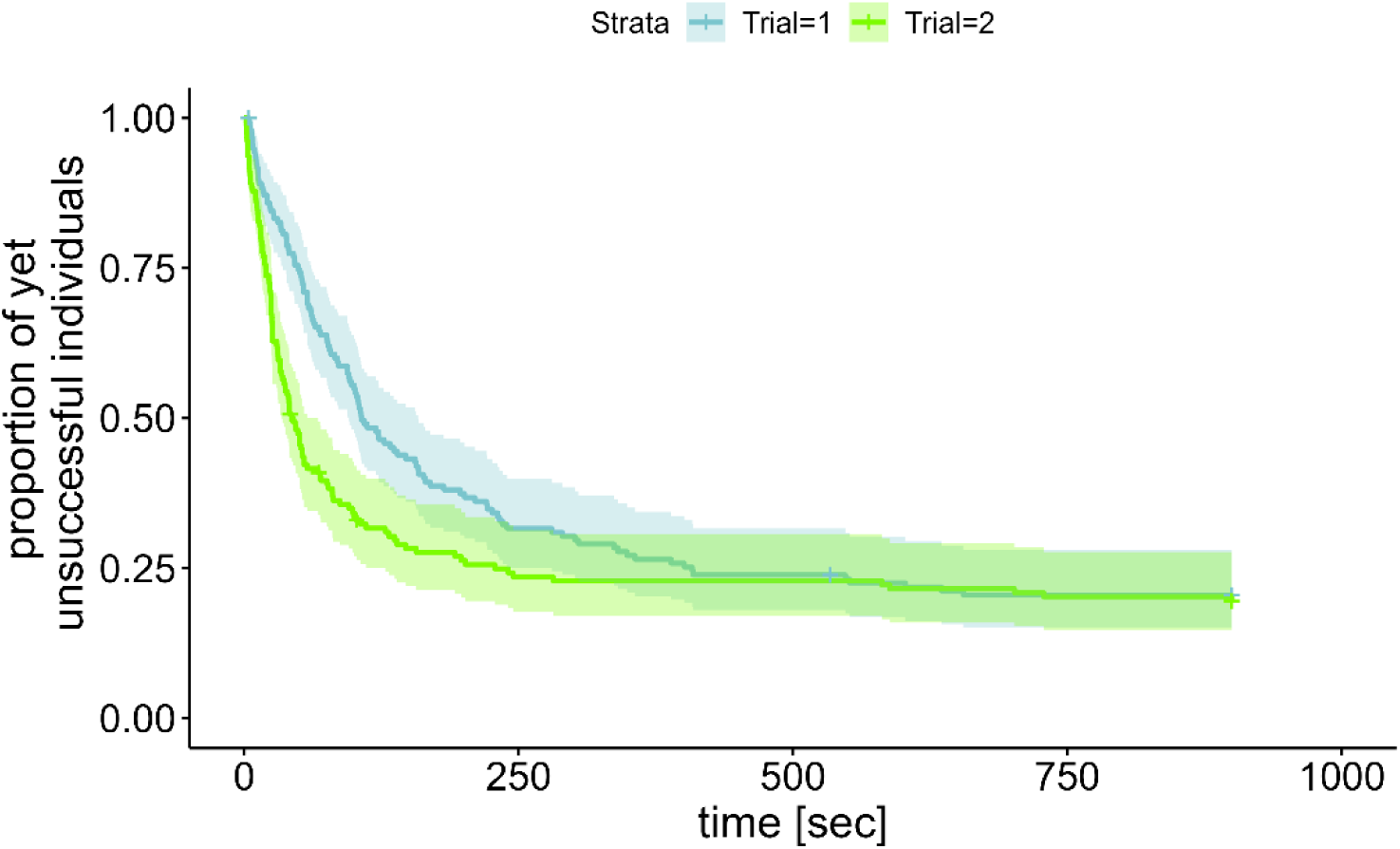
Kaplan-Meier plot comparing the latencies of finding the correct object in both trials of a session. Individuals found the correct object significantly faster during their second trial (green) than their first trial within the same session. See the online article for the color version of this figure.

Lastly, we examined whether individual differences like age, sex and status (i.e. being a helper vs a breeder) were associated with performance. Our model including only first interactions with the baited object as a correct choice revealed no effect of *age*, *sex* and *status* and was not significantly better than the null model (X2 (1) = 5.09, p = 0.16; see Table S4 in the Online Resource 1 for the full model summary).

For the DRT, we conducted a total of 543 trials. 14 subjects completed all 36 trials each, one female completed 27 trials, and one male completed 12 trials. The binomial GLMM revealed a significant positive intercept (GLMM: estimate = 0.57; standard error = 0.22, z-value = 2.5: *p* = 0.01), suggesting that across all conditions individuals performed significantly above chance level. Subsequently, we examined whether there was a difference in performance between the different delay conditions. The full model was better than the null model (X2 (1) = 34.96, p < 0.001) and revealed a significant linear decrease in performance with increasing delay (GLMM: estimate = -0.81; standard error = 0.15, z-value = -5.2: *p* < 0.001). Furthermore, the quadratic contrast of the delay condition also significantly influenced the performance (GLMM: estimate = 0.39; standard error = 0.15, z-value = 2.52: *p* = 0.01). Altogether the results suggest that the decline in performance is not constant with increasing delay but rather indicates a step decrease from the 0 seconds delay condition to the 15 seconds delay condition and almost no difference between the 15 seconds delay and the 30 seconds delay (Fig. 6). Furthermore, a subsequent analysis revealed that individuals did not perform above chance in either the 15-second or the 30-second delay scenario. Only when the delay was 0 seconds did animals perform better than expected by chance (GLMM: estimate = 1.3; standard error = 0.22, z-value = 2.5: *p* = 0.01).

**Fig. 6.**
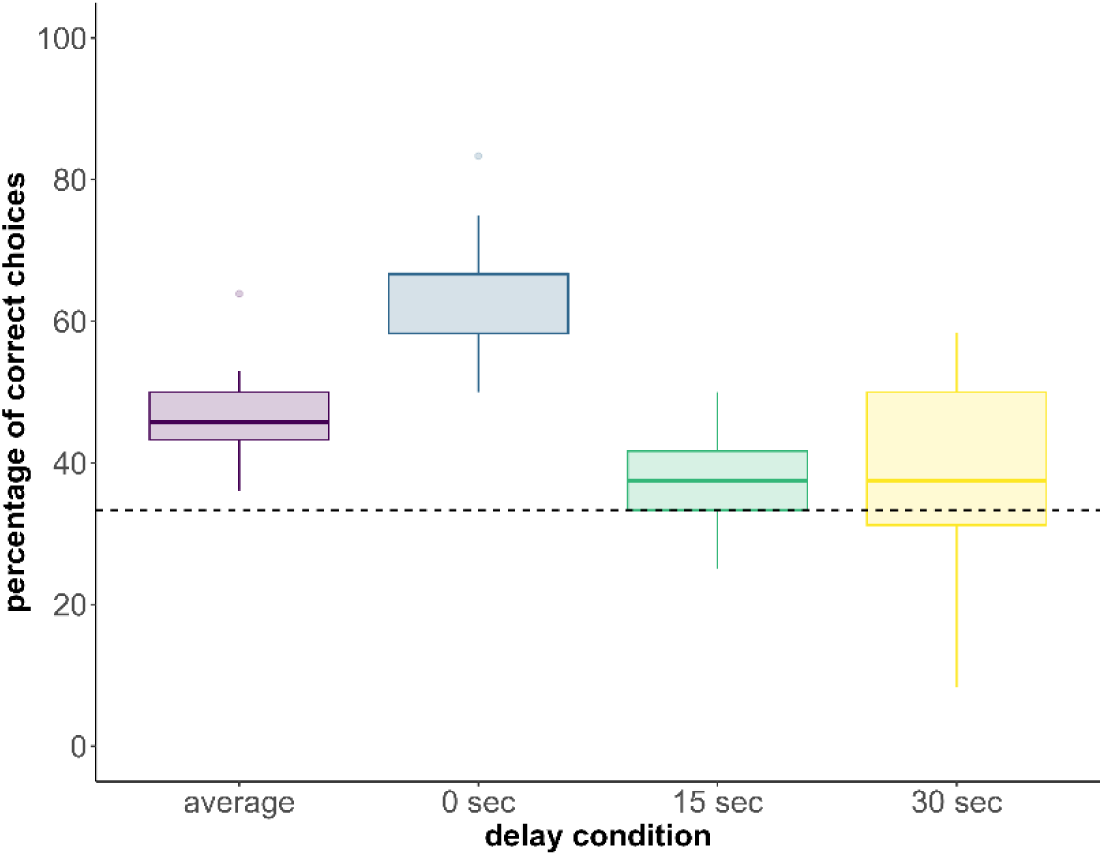
Boxplot of the percentages of correct choices separated into delay condition and average of all conditions. On average animals performed better than expected at chance-level (dotted line). However, this pattern was driven by the shortest delay as in the 0-second delay condition but not in the 15-second or 30-second condition, animals performed significantly better than by chance.

An important factor to consider for such tests is whether animals developed a side bias and preferably chose one cup more often than another regardless of the manipulation by the experimenter. We defined that animals have a side bias if they choose one cup more often than in one-third of all trials plus one standard deviation (in this case 12 + 1SD). The standard deviation was calculated by simulating random choices of 16 animals over 36 trials (12 and 27 trials for the individuals that did not complete all sessions respectively). Among the 16 tested animals, we identified 11 with a side bias (see Fig. S2 in the Online Resource 1). The marmosets that were considered biased chose the correct cup 46.1% (SD= 7.75%) of the time, while the non-biased chose the correct cup 47% (SD= 3%). Individuals with a side bias did not significantly perform worse than individuals without a side bias (GLMM: estimate = -0.04; standard error = 0.18, z-value = -0.21: *p* = 0.83).

The model containing *delay*, *age*, *sex,* and *status* as predictor variables and *family identity* as a random variable was significantly better than the null model (X2 (1) = 11.13, p = 0.02; see Table S5 in the Online Resource 1 for the full model summary). However, only delay had a significant effect on the performance, and neither age, sex nor status increased or decreased the model fit.

### Comparing the results of the 24h-DMT and the DRT

Ultimately, our objective was to examine a potential correlation between the number of correct choices observed in the 24h-DMT experiment and the DRT experiment (see Fig. S3 in the Online Resource 1). Using a Spearman’s correlation test, we found no correlation between the performances of the 24h-DMT experiment with the short delay DRT (ρ=−0.41; z = -1.61; p = 0.11), the middle delay (ρ=−0.21; z = 0.79; p = 0.43) or the long delay (ρ=−0.22; z = 0.84; p = 0.4). Furthermore, there was no correlation between the 24h-DMT experiment and the average performance of the DRT experiment (ρ=−0.12; z = -0.47; p = 0.64).

## Discussion

We examined memory capabilities of 16 marmoset monkeys using an open-foraging paradigm, the 24h-DMT, and the classical DRT and compared both results with each other. For the 24h-DMT, subjects were released individually into an unfamiliar environment containing up to 30 objects, with only one object containing food. Following a 24-hour delay, we assessed whether the monkeys retained the memory of the food’s location. In the second experiment we used the approach of the classical DRT. In the following we will first discuss the results of the 24h-DMT, followed by the results of the classical DRT and the comparison of both. Lastly, we will discuss why the classical DRT may not be a suitable test to examine memory and why even in captivity experiments with a more natural setup can provide a more accurate picture of the memory capacity of animals.

Our findings from the 24h-DMT align with previous research assessing spatial memory in marmoset monkeys. Marmosets perform well in open foraging tasks in both captivity (Easton et al., 2003; MacDonald et al., 1994; Platt et al., 1996; Vannuchi et al., 2020) and the wild (Abreu et al., 2020) and use heuristic rules to move between food locations (Xavier et al., 2024). In addition, like tamarins (Menzel & Juno, 1985), marmosets were able to remember food locations for 24 hours after a single trial of exploration. Although marmosets are omnivorous generalists, feeding on mobile prey like insects and small vertebrates, they also rely on (spatially and temporally) stable food sources like tree exudates (Souto et al., 2007). A substantial part of their diet comprises gum, and their home ranges are typically associated with the availability of gum trees. Their movement patterns often involve traveling between sleeping trees and gum trees as part of their daily activities (reviewed in Schiel & Souto, 2017). Hence, remembering the location of stable food sources like gum trees is important in the ecology of common marmosets. Our results also confirm that marmosets tend to rely on a *win-stay* strategy (MacDonald et al., 1994), meaning that they prefer to forage at locations where they have remembered being successful in the past. Food locations did not change between the first and the second trial of a session and marmosets tend to revisit the location where they found food during the last trial rather than during previous sessions. Their tendency to explore previously baited objects over novel objects first suggests a strong inclination toward the *win-stay* strategy. We also confirmed that marmosets would visit the baited object faster in their second trial of a session, corroborating that they remembered the location from the first session.

The results from the classical DRT follow the general trend of studies using this paradigm showing that an increase in the delay reduces the proportion of correct choices (Amici et al., 2010; Barth & Call, 2006; Call, 2001; Darusman et al., 2014; ManyPrimates et al., 2022; Schubiger et al., 2016). The marmosets performed rather poorly and performed at chance level at a delay of 15 seconds and 30 seconds. However, even in the no-delay condition which requires no memory, the performance of marmosets was below the average performance of other primate species in the same task (ManyPrimates et al., 2022). Given that the individuals were trained prior to the experiment, it is improbable that their performance is due to a lack of understanding of the task. Although we did not assess attention, it remains a likely explanation that the animals did not pay much attention to the experimenter and the task at hand and rather chose a side bias strategy which was on average successful in 33% of all trials. In fact, individuals with a side bias achieved the same number of rewards as those without. It has been shown that marmosets (and squirrel monkeys) pay more attention to such tasks when the probability of being successful is smaller (Schubiger et al., 2016). Similar results of such poor performances have been found in previous studies using marmosets as role models (Miles, 1957). In this classical study by Miles (1957), marmosets required over 1200 trials to perform at the same level as macaques in a 16-second delay DRT. Among other potential explanations, such as fundamental differences in their brain structure, it was speculated that marmosets exhibited poor attention to the task. In a similar study where the cues were given via a touch screen and after close to a hundred or more trials, four out of six marmosets made a correct decision after a 100-second delay (Yamazaki et al., 2016). Similar to our study, the marmoset tested was physically separated from other individuals but still in visual contact with at least one conspecific. Two key differences between our study and that of Yamazaki et al. (2016) may explain the variation in performance. First, the individuals tested in our study had less overall training and fewer experimental trials. Previous research suggests that marmosets improve their performance over hundreds of trials, eventually reaching nearly 90% accuracy (Miles, 1957; Yamazaki et al., 2016). An alternative explanation for improved memory could be that marmosets become well habituated to the test scenario, leading to reduced vigilance after hundreds of trials. This reduced vigilance may allow them to focus more attention on the stimuli. Second, in our study, clues were provided by an experimenter rather than automatically. Even minor variations in an experimenter’s behavior, despite standardized protocols, can influence an animal’s response.

Next, we examined the effect of individual traits on the overall performance of both experiments. Neither in the 24h-DMT, nor in the classical DRT could we pick up any factors that would explain individual differences. Against our prediction that performance would decline with age, we found no significant effect of age. Results on the effect of age on memory in non-human animals are mixed and sometimes reveal a significant decline (Darusman et al., 2014) or no effect (Flaim & Blaisdell, 2023; ManyPrimates et al., 2022). One possible explanation for this inconsistency is the influence of testing experience. Older individuals may compensate for their cognitive decline with testing experience. Although the here tested individuals have not been tested in either the 24h-DMT or the classical DRT (with the exception of Kapi and Vito (Schubiger et al., 2016); see Table S1 in Online Resource 1) before, experience from other cognitive tests may still facilitate their performance. Older individuals have participated in previous experiments (Brügger et al., 2021; Miss & Burkart, 2018; Schubiger et al., 2015; Sehner et al., 2022), whereas it was the first testing experience for younger animals. We also expected a better performance of females over males based on previous results from this colony (Schubiger et al., 2015). However, sex had no effect on the overall performance in both experiments, which may be due to the same reasoning as for age. The females in our dataset had less testing experience than the males. In addition, males are more aroused than females during tests in the presence of an experimenter (Schubiger et al., 2015), which in our case could only have had an effect on the second but not the first experiment.

Finally, we examined whether the performances of the two experiments correlate with each other. Overall, the performance in the DRT was much worse than in the 24h-DMT and we found no correlation between the tasks. Previous experiments have shown that information can be temporarily inaccessible and that memory performances after 24 hours and a sleeping period in between are better compared to shorter delays (Martin-Ordas & Call, 2011). This may be a contributing factor to the good performance during the 24h-DMT compared to the DRT but we argue that it cannot explain why the average performance in the 24h-DMT (80% correct choices) is better than the average performance in the no-delay condition of the DRT (65% correct choices). A second difference between the two experiments was the presence of an experimenter. Whereas the DRT required the presence of an experimenter at all times, the 24h-DMT was mostly performed in the absence of a person. Especially smaller species like marmosets can be influenced by the presence of an experimenter and attention to the task is impaired (Schubiger et al., 2015). Hence, marmosets in the DRT are likely to be more distracted than individuals in the 24h-DMT potentially decreasing performance. Lastly, the available information for the individuals was higher in the 24h-DMT compared to the DRT. In the former experiment, marmosets had access to spatial and visual information and could explore the locations of the hidden food reward longer than in the DRT. Again, spatial memory seems to decline more rapidly than other forms of memory (Rosenzweig & Barnes, 2003), meaning that performance is expected to be lower in the DRT compared to the 24h-DMT.

These results have implications for studies examining memory, in particular studies that aim at comparing between multiple species (e.g., Amici et al., 2010; ManyPrimates et al., 2022). The classical DRT represents one of the most repeated tests to assess memory, especially in primates (Amici et al., 2010; Barth & Call, 2006; Darusman et al., 2014; Herrmann et al., 2007; ManyPrimates et al., 2022; Schubiger et al., 2016). Despite its widespread application, our results raise doubts about its validity and call for adjustment of this paradigm, at least for marmosets and more likely for other species as well. Previous studies have shown that animals perform better when the experimenter secures the attention of the tested individual. In great apes, attention can be secured via both eye contact (Mulcahy & Call, 2009; Mulcahy & Suddendorf, 2011) and ostensive signals (Kano et al., 2018). However, depending on the social structure, some species may react threatened to eye contact or do not respond to ostensive signals (Harrod et al., 2020). Another possibility to achieve increased attentiveness and thus more accurate measurement of memory performance is to decrease the probability of success in such tests (Schubiger et al., 2016). Common marmosets and squirrel monkeys paid more attention to the test when the potential costs of not paying attention were higher. However, the overall performance of non-human primates in these classical DRTs remains rather poor or moderate at best, showing that even great apes have problems locating the right container after only a couple of seconds (ManyPrimates et al., 2022).

An alternative and less artificial way of assessing memory is to use open foraging paradigms. Compared to the classical DRT, performances in these open-foraging paradigms are less affected by delay and correct choices remain above chance level in conditions of shorter (Platt et al., 1996)- and longer delay (Abreu et al., 2020; Menzel & Juno, 1985; but see Platt et al., 1996). Additionally, open foraging paradigms can be conducted in a way that the presence of an experimenter is unnecessary and without the necessity to train animals on an automatic apparatus, thus avoiding potential interferences of different individuals toward one or multiple experimenters (Schubiger et al., 2015). Open-foraging paradigms also seem to circumvent the problem of motivational biases mentioned by Schubiger et al. (2016) since animals perform well above chance with only a few items to choose from (Abreu et al., 2020) and multiple items as seen here and in previous results (Menzel & Juno, 1985). Although the chance level cumulatively decreased with increasing sessions and was ultimately much lower than in the classical DRT, performance in the 24h-DMT did not decrease. In addition, individuals can collect and memorize information rather independently as they would under natural conditions. This undermines the capacity to memorize food locations in common marmosets if tested in a paradigm that has some ecological relevance. Marmoset diet heavily relies on exudates (Schiel & Souto, 2017) and marmosets have to remember where these food sources are located. Hence, like in the 24h-DMT paradigm, they freely explore their environment and memorize spatial and visual information about food locations. Finally, unlike the classical DRT, open-foraging paradigms can be applied to wild populations (Abreu et al., 2020), thus allowing us to test memory under natural conditions.

## Acknowledgments

We are grateful to Gisep Bazzell for taking care of the animals and assisting with their handling. Thanks to Eloisa Martins for her support in creating the images of the apparatuses and test enclosure. We are grateful to Rodolph Schlaepfer for his help with the camera setup. A previous version of this manuscript greatly benefited from the comments of two anonymous reviewers. This project was supported by an SNF Grant (grant number 31003A_172979) to Judith Burkart and an SNF Postdoc.Mobility Grant (grant number P500PB_217864/1) to Sandro Sehner.

## Declarations

### Data availability

All data needed to evaluate the conclusions in the paper are present on OSF (DOI 10.17605/OSF.IO/3V6SZ).

### Ethical approval

All the experiments were in accordance with the Swiss legislation and licensed by the Kantonales Veterinäramt Zürich (license number: ZH232/19, degree of severity: 0).

