## Supplemental material for "Common marmosets (*Callithrix jacchus*) excel in a one-trial spatial memory test, yet perform poorly in a classical memory task"

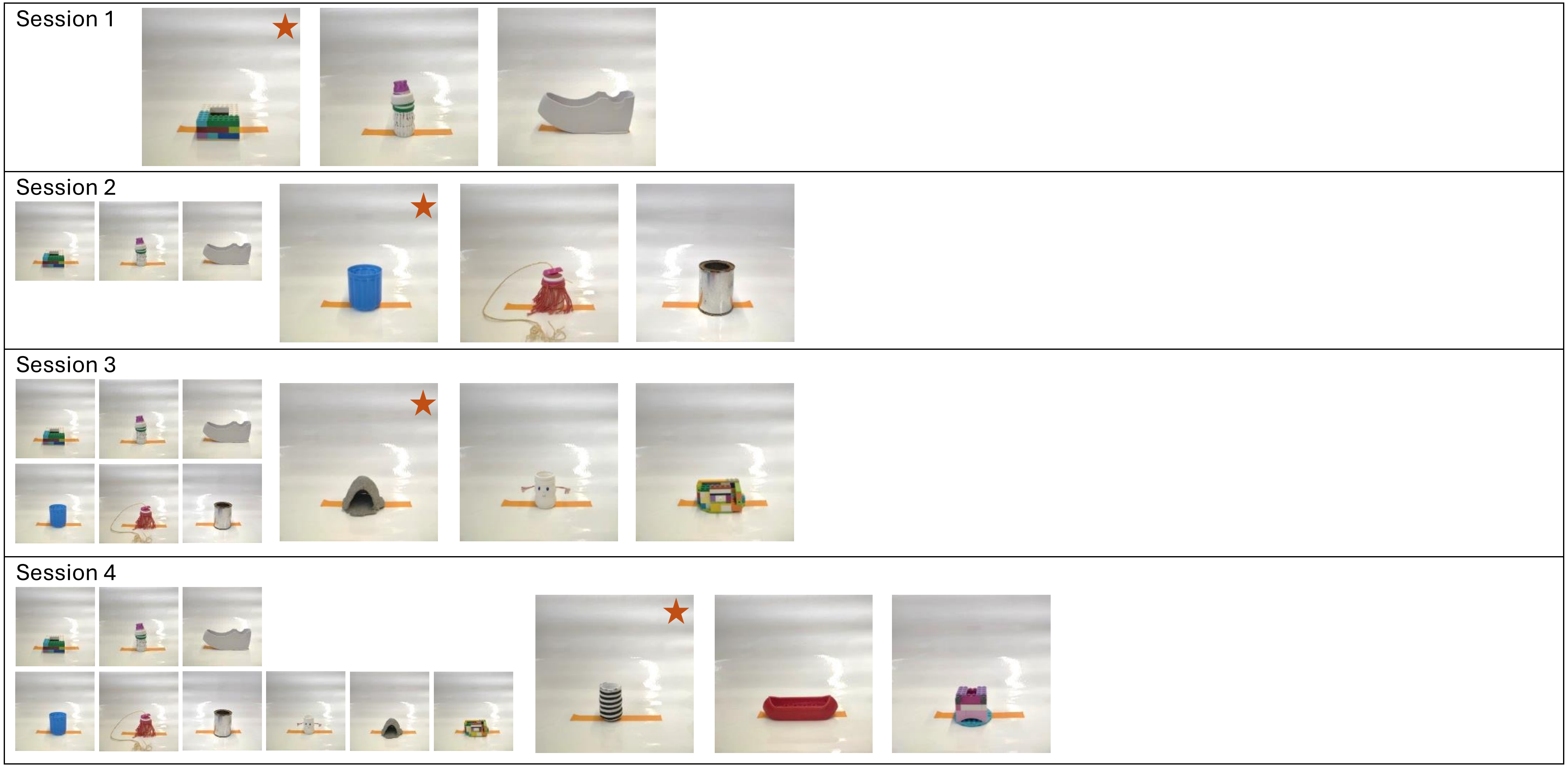
Session 1
Session 2
Session 3
Session 4

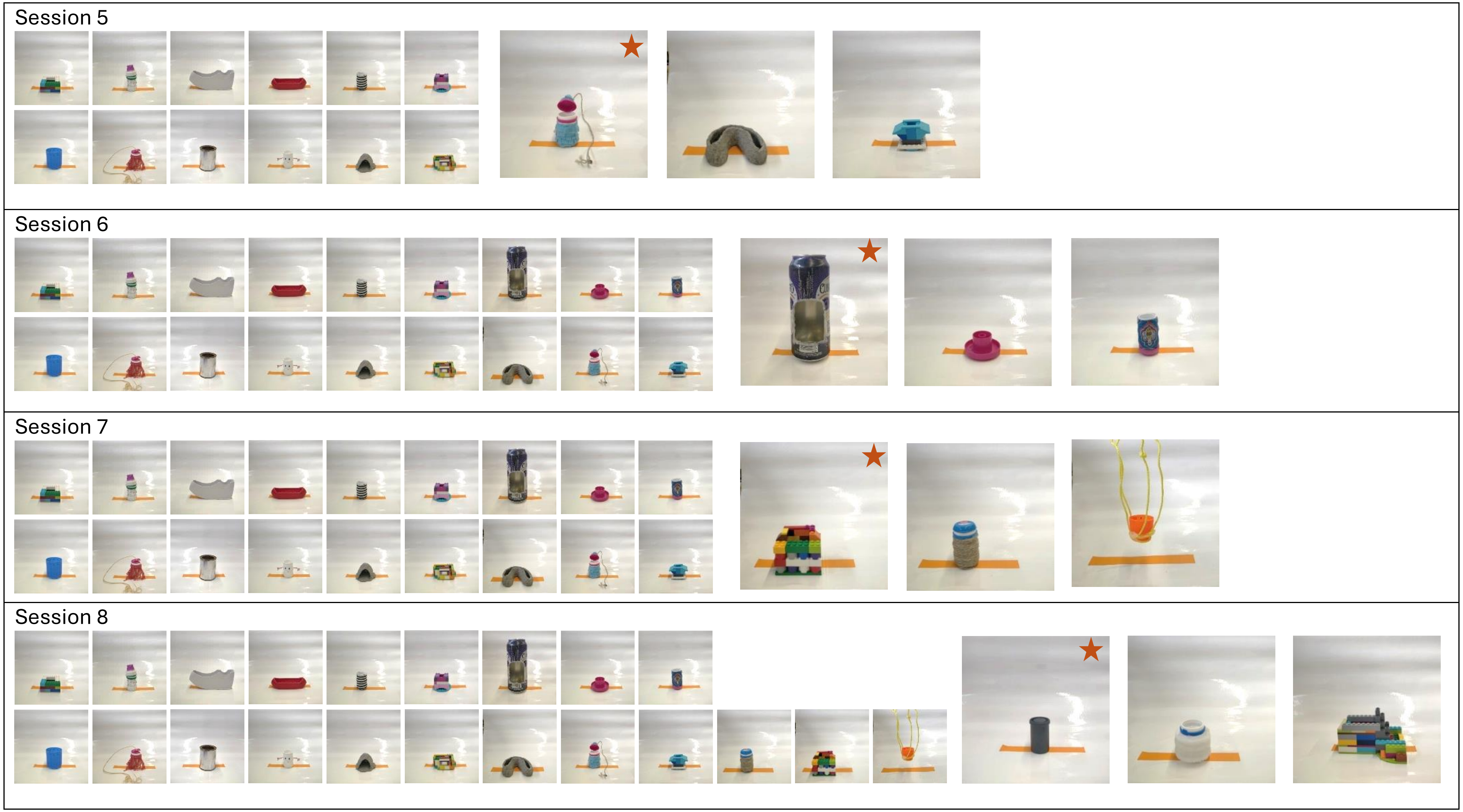
Session 5
Session 6
Session 7
Session 8

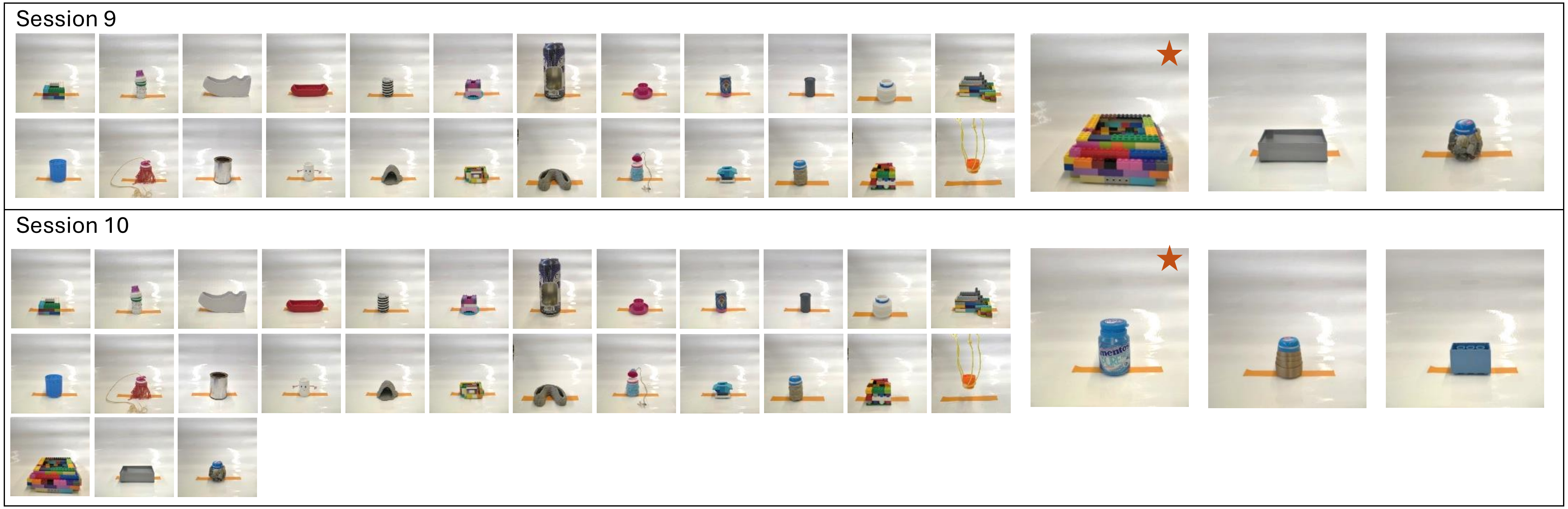


**Fig. S1**. Images of all objects that were used for the 24h delay experiment. The larger images on the right represent the three objects that were added in that specific session. Images marked with a red star at the top right corner indicate the this object was baited during this session. Previously introduced objects remain in the enclosure and thus the number of objects accumulates from three (Session 1) to 30 (Session 10). However, only one object is baited during each trial. The orange stripe of tape is added as a reference and is 15 cm long, but was not added in the enclosure.

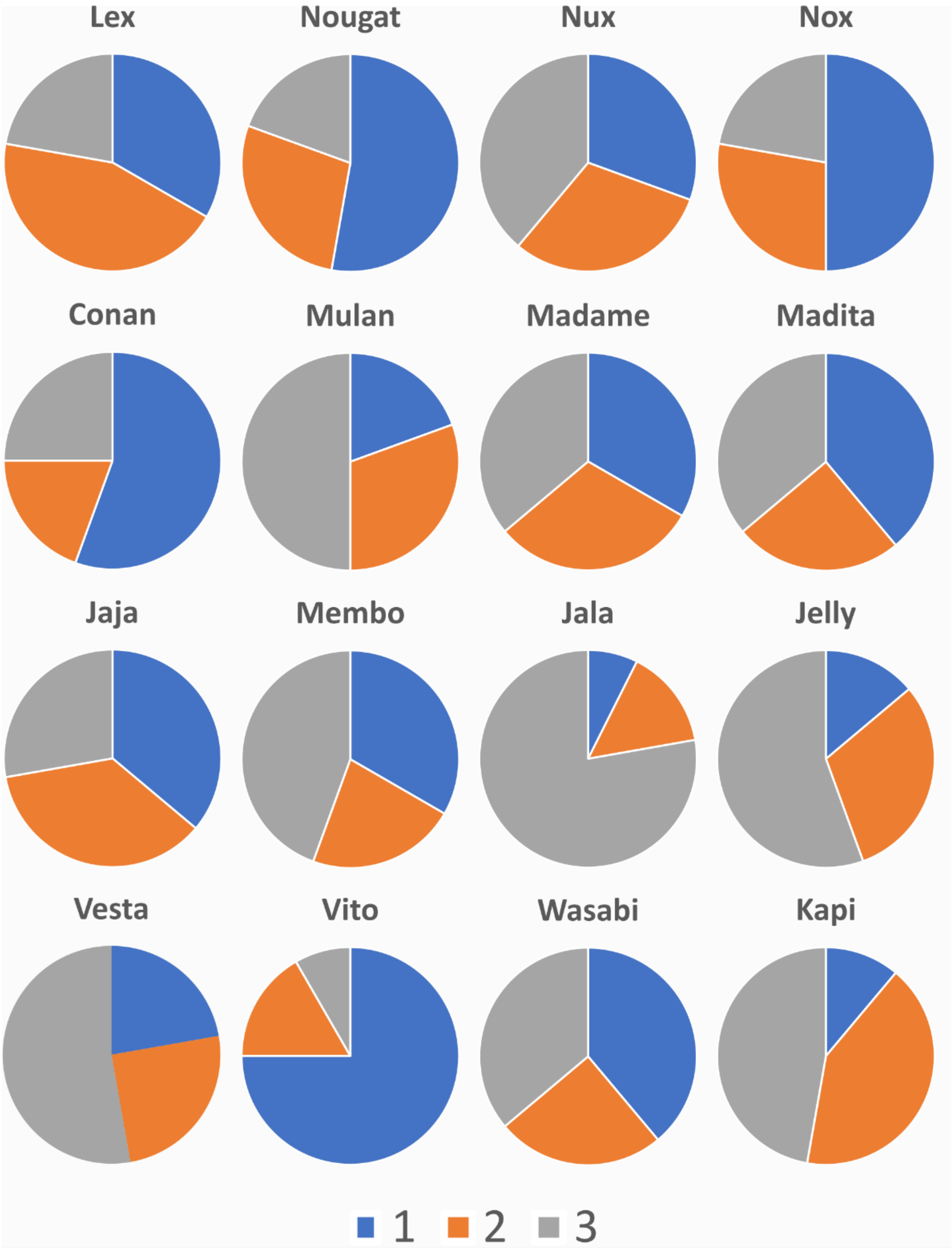


**Fig. S2.** Depiction of the choices of each animal based on the side they choose. Position 1 (blue) refers to the object on the right-hand side from the animal's perspective; Position 2 (orange) refers to the object in the central position; Position 3 (grey) refers to the object on the left-hand side from the animal's perspective. We defined that animals had a side bias if they chose one position more often than in one-third of all trials plus one standard deviation of a set of simulated data (in this case 12 + ≈ 3). Eleven animals were considered to have a side bias (all but Madame, Madita, Nux, Wasabi, and Jaja).

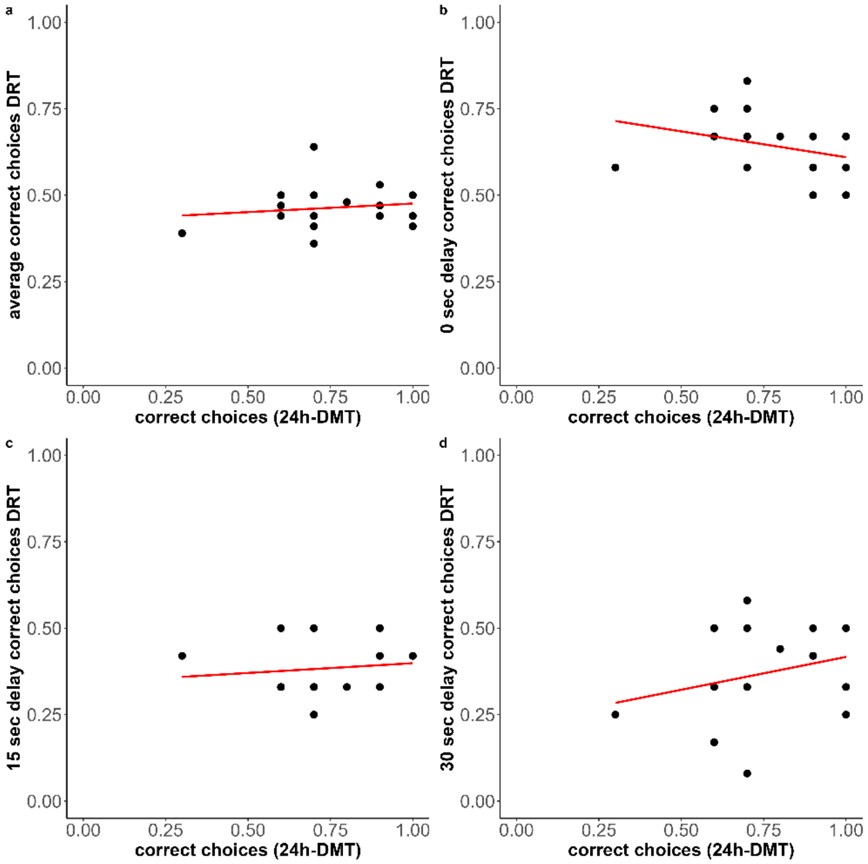


**Fig. S3.** Correlation plot between the proportion of correct choices of each individual during the 24h-DMT and the a) average proportion of correct choices in the DRT; b) the proportion of correct choices in the zero delay condition in the DRT; c) the proportion of correct choices in the middle delay condition in the DRT; and d) the proportion of correct choices in the long delay condition in the DRT. None of these correlations were significant.

**Table S1.** Summary of all individuals participating in the 24h-delay experiment. Sex is categorized into females (F) and males (M), while status is distinguished between breeders (B) and helpers (H), resulting in four distinct categories. The number of correct first interactions refers to the second trial, where animals had to remember which object contained the food. The last column refers to the average difference in latency when interacting with the currently baited object between the first and second trials. This average only considers sessions where the individual found the correct object in the first trial.

| Family | Individual | Sex and status | Age [years] | Number of correct first interactions | Average difference of interacting with the correct object on the first and second trial [sec] |
|---|---|---|---|---|---|
| Mibbas | Conan | MB | 6 | 7 | 103.84 |
| | Mulan | FH | 2.5 | 9 | 93.16 |
| | Madame | FH | 1.5 | 9 | 55.30 |
| | Madita | FH | 1.5 | 10 | 73.72 |
| Jajas | Jaja | FB | 11 | 8 | 224.36 |
| | Membo | MB | 11 | 7 | 49.18 |
| | Jala | FH | 5.5 | 7 | 73.3 |
| | Jelly | MH | 4 | 9 | 27.8 |
| Ninas | Lex | MB | 14 | 6 | 146.8 |
| | Nougat | FH | 5 | 7 | 183.71 |
| | Nux | FH | 4 | 10 | 144.06 |
| | Nox | MH | 4 | 10 | 87.97 |
| Vestas | Vesta | FB | 16 | 6 | 35.55 |
| | Vito | MB | 14 | 6 | 243.63 |
| Wasabis | Wasabi | FB | 3.5 | 7 | 103.52 |
| | Kapi | MB | 18 | 3 | 223.39 |

**Table S2.** Exemplary scoresheet of the ManyPrimates-Short-Term Memory experiment. ManyPrimates provided the sequences of conditions and hiding locations.

| **Subject:** | | | | **Experimenter** | | |
|---|---|---|---|---|---|---|
| **Date** | **Block** | **Trial** | **Condition** | **Hiding Location** | **Pick** | **Correct** |
| | 1 | 1 | Long | 3 | | |
| | 1 | 2 | Long | 2 | | |
| | 1 | 3 | Long | 1 | | |
| | 2 | 4 | Medium | 1 | | |
| | 2 | 5 | Medium | 2 | | |
| | 2 | 6 | Medium | 3 | | |
| | 3 | 7 | Short | 1 | | |
| | 3 | 8 | Short | 2 | | |
| | 3 | 9 | Short | 3 | | |

**Table S3.** Table with polynomial contrasts for correct choices comparing earlier with later sessions. None of the contrasts revealed significance, indicating that animals made as many correct choices in earlier sessions as they did in later sessions, despite an increasing number of objects.

| Types of polynomial contrasts | p-values |
|---|---|
| L | 0.98 |
| Q | 0.97 |
| C | 0.97 |
| ^4 | 0.97 |
| ^5 | 0.97 |
| ^6 | 0.98 |
| ^7 | 0.97 |
| ^8 | 0.97 |
| ^9 | 0.97 |

**Table S4.** Summary of the full model of correct choices (interacting with the baited object first) in the second trial of the 24-hour delay experiment, including *age*, *sex,* and *status* as predictor effects and *session number* and *individual identity nested in family identity* as random effects. Bold values indicate $p \leq 0.05$.

| **Fixed factors** | **Odds ratio** | **95% CI** | **Estimate** | **SE** | **Z** | **p** |
|---|---|---|---|---|---|---|
| *Intercept* | 0.91 | 0.33 – 2.51 | -0.09 | 0.51 | -0.17 | 0.86 |
| *Age* | 0.97 | 0.89 – 1.05 | -0.03 | 0.04 | -0.84 | 0.40 |
| *Sex (M)* | 1.26 | 0.55 – 2.9 | 0.23 | 0.42 | -0.55 | 0.58 |
| *Status (H)* | 1.98 | 0.85 – 4.57 | 0.68 | 0.43 | 1.59 | 0.11 |

**Table S5.** Summary of the full model of correct choices (finding the hidden food item) in delayed response task, including *age*, *sex,* and *status*, and *delay* as predictor effects and *individual identity nested in family identity* as random effect. Bold values indicate $p \leq 0.05$.

| **Fixed factors** | **Odds ratio** | **95% CI** | **Estimate** | **SE** | **Z** | **p** |
|---|---|---|---|---|---|---|
| *Intercept* | 0.91 | 0.59 – 1.41 | -0.09 | 0.22 | -0.39 | 0.7 |
| *Age* | 0.99 | 0.96 – 1.03 | -0.01 | 0.02 | -0.37 | 0.71 |
| *Sex (m)* | 1.14 | 0.77 – 1.68 | 0.2 | 0.2 | 0.65 | 0.52 |
| *Status (h)* | 0.91 | 0.62 – 1.32 | 0.19 | 0.19 | -0.51 | 0.61 |
| *Delay (linear)* | 0.44 | 0.32 – 0.60 | -0.81 | 0.15 | -5.23 | **< 0.001** |
| *Delay (quadratic)* | 1.47 | 1.09 – 1.99 | 0.38 | 0.15 | 2.53 | **0.01** |
